# Social threat alters the behavioral structure of social motivation and reshapes functional brain connectivity

**DOI:** 10.1101/2024.06.17.599379

**Authors:** Geronimo Velazquez-Hernandez, Noah W. Miller, Vincent R. Curtis, Carla M. Rivera-Pacheco, Sarah M. Lowe, Sheryl S. Moy, Anthony S. Zannas, Nicolas C. Pégard, Anthony Burgos-Robles, Jose Rodriguez-Romaguera

## Abstract

Traumatic social experiences redefine socially motivated behaviors to enhance safety and survival. Although many brain regions have been implicated in signaling a social threat, the mechanisms by which global neural networks regulate such motivated behaviors remain unclear. To address this issue, we first combined traditional and modern behavioral tracking techniques in mice to assess both approach and avoidance, as well as sub-second behavioral changes, during a social threat learning task. We were able to identify previously undescribed body and tail movements during social threat learning and recognition that demonstrate unique alterations into the behavioral structure of social motivation. We then utilized inter-regional correlation analysis of brain activity after a mouse recognizes a social threat to explore functional communication amongst brain regions implicated in social motivation. Broad brain activity changes were observed within the nucleus accumbens, the paraventricular thalamus, the ventromedial hypothalamus, and the nucleus of reuniens. Inter-regional correlation analysis revealed a reshaping of the functional connectivity across the brain when mice recognize a social threat. Altogether, these findings suggest that reshaping of functional brain connectivity may be necessary to alter the behavioral structure of social motivation when a social threat is encountered.

## Introduction

Survival in the real world requires constant regulation of behavioral responses to salient stimuli. Previous experiences are critical to adapt behavioral responses and enhance survival [1–3]. These adaptive responses are often classified into bimodal motivated behavioral responses characterized by either approach toward a safe stimulus or avoidance away from a threatening stimulus [4–8]. These bimodal responses also exist towards social stimuli, and can help us understand if a social stimulus is perceived as safe or threatening. However, the repertoire of behaviors performed during social encounters is far richer than simple approach and avoidance [9]. Studying social behaviors provides insights into how an individual interacts within its broader societal context, their inclination toward forming social bonds, and their aversion to unfavorable social interactions [10]. Therefore, understanding the complex nature of social behaviors is essential for delineating maladaptive behaviors associated with psychiatric disorders linked to social deficits, such as social anxiety and autism spectrum disorders.

Social experiences create memories that enable individuals to categorize subsequent social encounters as either threatening or safe, thereby eliciting appropriate behaviors to achieve specific goals [11–16]. Prior studies have shown that multiple brain regions signal distinct socially motivated behaviors; for example, the ventromedial hypothalamus (VMH), medial amygdala (MeA), and the bed nucleus of stria terminalis (BNST) are implicated in promoting aggression and defensive responses [17–22], while the nucleus accumbens (NAc), medial preoptic area (mPOA), and ventral tegmental area (VTA), among others, facilitate social interactions [10,16,23–29]. In addition, these regions have also been shown to regulate stress, aversion, reward, and decision-making [7,30–36]. Evaluating and manipulating individual brain regions in isolation allows us to study their functional role ina specific behavioral output [37–40]. However, we must also understand how global activity dynamics among them orchestrates such behaviors [41,42].

To better understand the full repertoire of behavioral change in the presence of a social threat, we first employed a combination of classic and recently developed behavioral analysis techniques that allow sub-second analysis of behavior during social learning tasks. In this task, mice learn to associate another mouse as either threatening (social contact paired with foot-shock) or safe (no foot-shock). Using both single-point tracking (one point on mouse body) and multi-point tracking (key points across body and tail) with motion sequencing, we demonstrate that social threat recognition alters the behavioral structure of social motivation. Furthermore, we employ an inter-regional correlation analysis approach of cFos+ cell counts, and we find that recognizing a social stimulus as a threat reshapes inter-regional correlations in cFos+ cell count across regions that regulate stress, aversion, reward, decision-making, and social motivation. Taken together, our findings highlight how reshaping of functional brain connectivity may orchestrate the behavioral structure of social motivation.

## Materials and Methods

### Subjects

Adult male and female C57BL/6J mice (Jackson Laboratories) weighing between 22-30g at the beginning of the experiment were used for the study. All procedures were conducted in accordance with the Guide of the Care and Use of Laboratory Animals, as adopted by the National Institute of Health, and with the approval of the Institutional Animal Care and Use Committee from the University of North Carolina at Chapel Hill.

### Social threat learning

To evaluate behavioral responses occurring from threatening and safe social interactions, we used modular test chambers (ENV-007-VP, Med Associates, VT, USA) (Figure 1A) in which social contact was paired with foot-shock (social threat) or not (social safe) [43]. To house the stimulus mouse, we built a compact social chamber (6x6x9 cm); in which the chamber was aligned 2 cm below the floor grids, compelling the stimulus mouse to rear in order to engage with the experimental mouse. To ensure alignment of social contact with a foot-shock (1 second, 0.3 mA), we used two photobeam systems that had to be simultaneously triggered on either side of an interaction window. We considered social contact to occur when the photobeams separating the two animals were broken (78% of mice completely avoided after the first trial).

**Figure 1.**
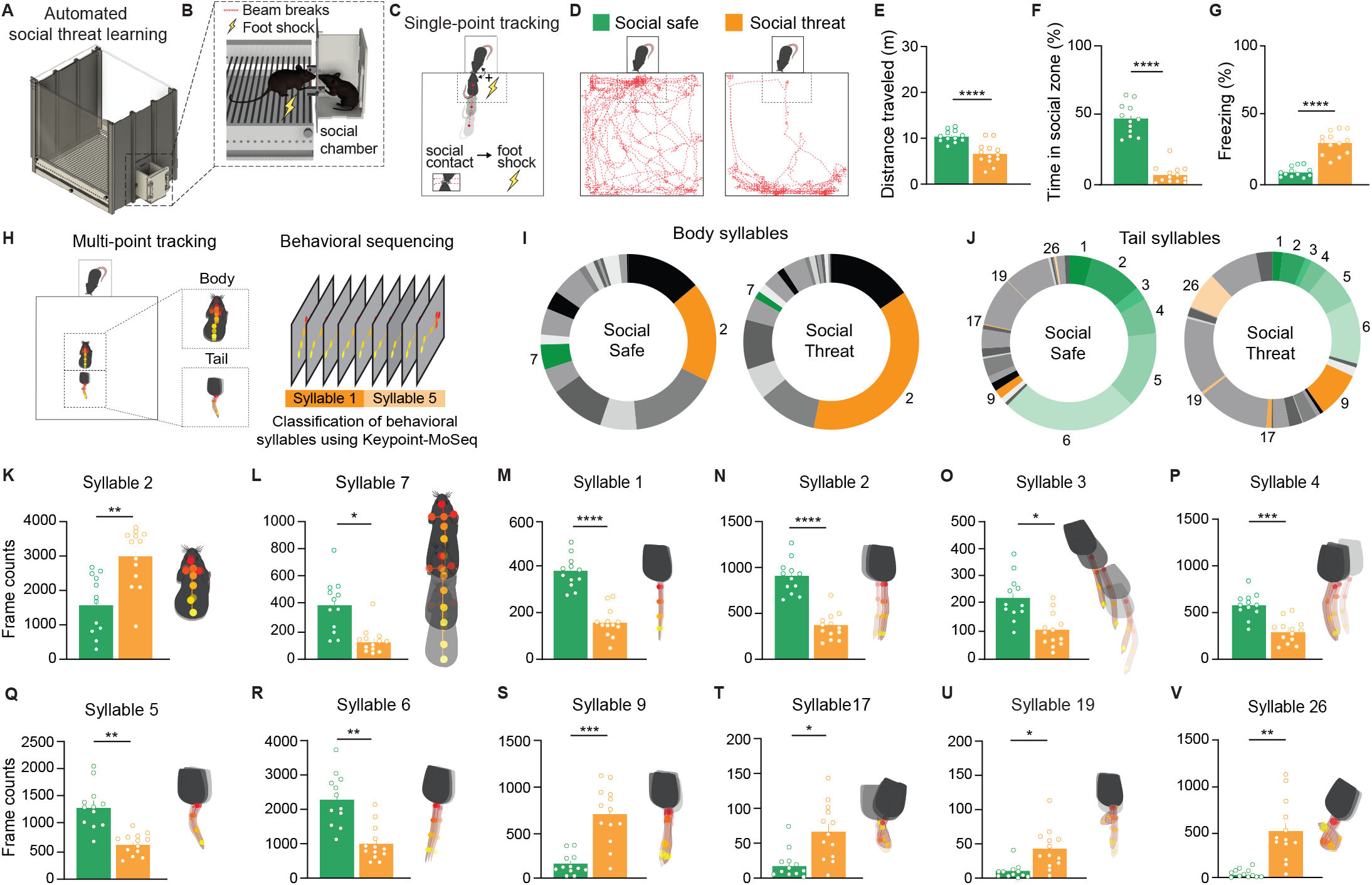
Social threat learning elicits multiple behavioral syllables in mice that are related to defensive behaviors. **(A)** Schematic representation of the automated social threat learning chamber. **(B)** Lateral depiction of social threat learning, illustrating how social interactions interrupt photobeams, triggering foot-shock delivery. **(C)** Cartoon representation of single-point tracking analysis during SFC using the AnyMaze software. **(D)** Representative traces from single-point tracking, highlighting behavior navigation of a social safe and social threat mouse. **(E)** Distance traveled decreased in the social threat group (t(23) = 4.88, p = 0.0001). **(F)** Social approach decreased in the social threat but not the social safe group, indicated by time in the social zone (t(23) = 10.95, p = 0.0001). **(G)** Freezing increased in the social threat but not the social safe group (t(23) = 7.81, p = 0.0001). **(H)** For comprehensive behavioral analysis, body and tail movements were analyzed separately during SFC using multipoint tracking analysis (left), enabling the classification of behavioral syllables given through unsupervised machine learning (Keypoint-MoSeq) (right). **(I)** Nineteen distinct syllables were identified for body movements, and **(J)** twenty-eight for tail movements. Gradient colors represent syllables expressed more in social safe mice (green) or social threat mice (orange). **(K-L)** Comparison of body frame counts between groups shows a significant difference on syllable 2, p = 0.003, and syllable 7, p = 0.005. **(M-V)** Comparison of tail frame counts between groups shows significant differences on multiple syllables (F(27, 621) = 14.16, p = 0.0001: **(M)** syllable 1 p = 0.0001, **(N)** syllable 2 p = 0.0001, **(O)** syllable 3 p = 0.01, **(P)** syllable 4 p = 0.002, **(Q)** syllable 5 p = 0.001, **(R)** syllable 6 p = 0.002, **(S)** syllable 9 p = 0.0007, **(T)** syllable 17 p = 0.04, **(U)** syllable 19 p = 0.04, **(V)** syllable 26 p = 0.008. **(K-V)** Diagram on right: recreation of the multi-point tracking analysis sequence. Social safe, n= 12; social threat, n= 13.

### Three-chamber social interaction

To evaluate social behavioral responses after social threat learning, we exposed mice to a classic three-chamber social assay (60 cm x 35 cm) 24 hours following social threat learning [50].

### Behavioral analysis

For single-point tracking we used AnyMaze Software (Stoelting Co., IL, USA). For the tracking of key points along the body and tail, we used DeepLabCut [44]. The DeepLabCut model was trained by labeling 20 representative frames from 25, 10-minute videos (500 frames total). Body (6 points) and tail (4 points) labels were distributed evenly across respective regions. For the extraction of sub-second behaviors, we used Keypoint-MoSeq [45], a behavioral clustering algorithm that finds statistically significant behavioral syllables using keypoints obtained from DeepLabCut. Keypoint-MoSeq employs a probabilistic time-series model to identify behavioral syllables that exhibit consistent patterns of occurrence.

### Immunohistochemistry

To investigate the potential brain regions involved in modulating social recognition, we forced social interaction on day 2. For this purpose, we placed the experimental and stimulus mice in a chamber separated by a barrier with two windows and let them interact for 10 minutes. 90 minutes after interaction, mice were anesthetized with isoflurane (drop method), transcardially perfused, and cFos immunohistochemistry was performed (see Extended Methods).

### Cell counting and analysis

We quantified the number of cFos+ nuclei in regions related to either defensive and/or social behaviors. Seventeen brain tissue sections spanning the anterior-posterior axis of each of these regions were processed for analysis. Data collected from separate brain hemispheres were counted as independent samples and data from matching regions across sections were averaged (n=4 mice sampled per condition, totaling a max of n=8 hemispheres per condition used for analysis). Images of brain sections were registered to the Allen Mouse Brain Atlas V3 using Aligning Big Brains & Atlases (ABBA), and quantification of total cFos+ cells counts was performed using QuPath v04.4 [46].

### Statistical analysis

Groups were compared by using, when appropriate, unpaired Student’s two-tailed t-tests, one-way, or repeated-measures analysis of variance (ANOVA) followed by *post-hoc* Sidak or Tukey Least Significant Difference test using Prism (GraphPad, La Jolla, CA). Inter-regional correlation analyses were performed by calculating Pearson’s correlation coefficients for regional pairs using R-Studio. For all analyses, alpha was set at p<0.05.

## Results

Traditional behavioral tracking methodologies have explored socially motivated behavioral responses through the evaluation of simple avoidance and approach responses. Many studies have also focused on unique experimenter-defined social behaviors such as aggression, social attention, or stretching [47,48]. These studies often require careful visual observation and biased quantification by an expert. More recently, the use of unsupervised machine learning techniques for unbiased behavioral analysis has enabled a more thorough investigation into entire behavioral sequences with sub-second precision [45,49]. In the present study, we employed an automated social threat learning task that pairs social contact with foot-shock. We use both traditional single-point tracking (AnyMaze), as well as multi-point tracking techniques to perform behavioral sequencing and extract behavioral syllables (Keypoint-MoSeq) [45,49]. To quantify global brain activity changes, we employ automated brain atlas registration and cell detection techniques (ABBA and QuPath) to quantify cFos+ expression patterns and perform inter-regional correlation analysis across multiple brain regions of interest.

### Social threat learning alters sub-second behavioral sequences

To evaluate social threat learning, we automated a behavioral task that involves administering a foot-shock upon social contact [21,43] (Figure 1A-B). An experimental mouse was placed in a testing chamber, and, following a baseline period, a stimulus mouse was introduced into an attached stimulus chamber that allowed social contact via an interaction window. For the “social threat” group, the mouse received a foot-shock every time it made social contact with the stimulus mouse (Figure 1C). The same experimental design was employed with a “social safe” group, in which mice were allowed social contact with stimulus mice but did not receive a foot-shock. To initially assess behavioral differences between the groups, we collected classical behavioral outputs using single-point tracking (ANY-maze; Stoelting Co., IL, USA). No differences between groups were observed during the initial baseline period, when mice were exposed to the chamber in the absence of a social stimulus (Supplementary Figure 1B-C). During the social threat learning period, the social threat group exhibited a decrease in distance traveled (Figure 1D-E), time spent in the social zone (Figure 1F), and increased freezing (Figure 1G), in comparison to the social safe group.

To uncover novel behavioral syllables associated with these types of social interactions, we implemented Keypoint-MoSeq, an unsupervised machine learning model capable of extracting statistically relevant sub-second behaviors from keypoint data collected using DeepLabCut (Figure 1H). We trained separate models with key points from the body and the tail. Following training, the model returned a total of 19 different behavioral syllables for the body and 29 for the tail (Figure 1I-J). During the initial baseline period, we found no differences between the groups for both the body and tail (Supplementary Figure 1E). However, during social threat learning, we found significant differences across multiple behavioral syllables for both the body and tail. We found two distinct syllables for the body (Figure 1 K-L), and 10 distinct syllables for the tail (Figure 1 M-V). To illustrate the behaviors from the body and tail related to the syllables, we generated illustrations for each significant syllable (Figure 1K-V). When assessing potential distinctions between males and females across significant syllables, we observed that social threat learning induced higher occurrence of two syllables in females over males in body syllable 9 and tail syllable 17 (Supplementary Figure 2K-L). Overall, these findings demonstrate that when mice have a social interaction that is paired with foot-shock, they exhibit classic defensive responses such as avoidance and freezing, as well as an entire repertoire of sub-second behavioral sequences in both the body and the tail. Furthermore, it’s important to note that some of these behavioral changes may be due to the painful experience of the foot-shock, outside of the aversive nature of the social interaction experienced.

### Social threat learning alters sub-second behavioral sequences

To evaluate social threat recognition, we exposed mice to a three-chamber social interaction assay the day after either social threat or safety learning. Using single-point tracking (Figure 2A) [50], we found no differences in distance traveled between the social threat and social safe groups (Figure 2C), but we found that the social threat group avoided exploring the social side, while the social safe group preferred it (Figure 2D, Supplementary Figure 4A-B). Freezing responses increased in the social threat group, as compared to the social safe group (Figure 2E). These results suggest that social threat learning creates a strong social memory that allows animals to recognize a social threat and promote defensive responses.

**Figure 2.**
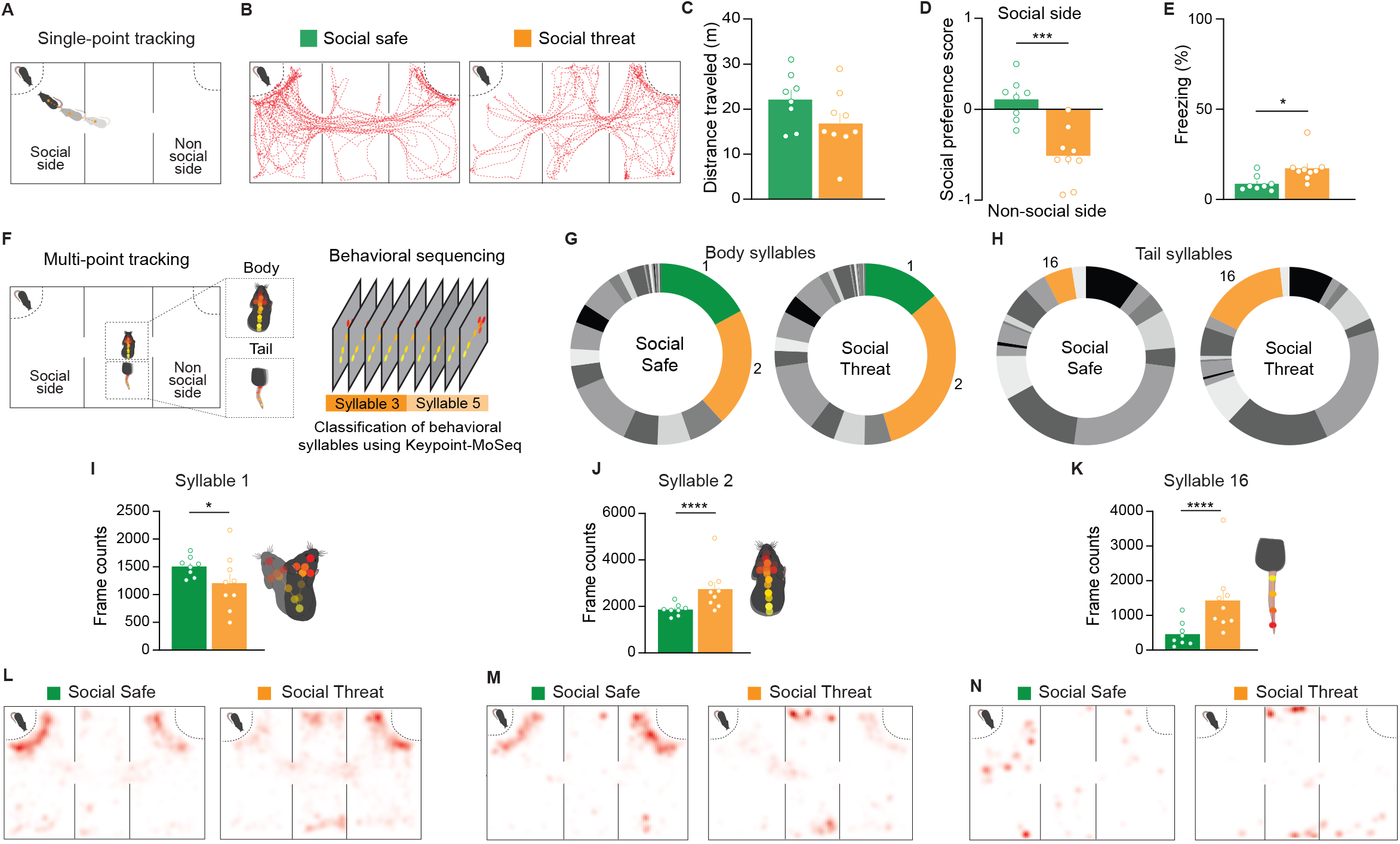
Social threat recognition induces unique behavioral syllables. **(A)** Schematic representation of three-chamber social interaction assay and single-point tracking analysis using the AnyMaze software. **(B)** Representative traces from single-point tracking, highlighting behavior navigation of a social safe and social threat mouse. **(C)** Motricity is not affected by social threat memory, as indicated by similar distance traveled by social safe and social threat groups (t(15) = 1.45, p = 0.16). **(D)** The social threat group avoids the social side, exploring the non-social side more, while the social safe group explores the social side more than the non-social side (t(15) = 4.73, p = 0.003). **(E)** Social threat groups showed an increase in freezing during social recognition (t(15) = 2.66, p = 0.01). **(F)** For a comprehensive behavioral analysis, body and tail movements were analyzed separately during SFC using multipoint tracking analysis (left), enabling the classification of behavioral syllables given through a statistical model in an unsupervised way (Keypoint-MoSeq) (right). **(G)** Twenty-three distinct syllables were identified for body movements and **(H)** seventeen for tail movements. The green color represents syllables expressed more in social safe mice, and the orange color represents syllables expressed more in the social-threat mice. **(I-J)** Comparison of body frame counts between groups only on syllables with a statistical difference (F(22, 105) = 4.85, p = 0.0001, posthoc **(I)** syllable 1, p = 0.04, **(J)** syllable 2, p = 0.001). **(K)** Comparison of tail frame counts between groups only on syllables with statistical difference (F(16, 255) = 3.82, p = 0.001, posthoc syllable 16, p = 0.0001). **(L-N)** Heatmaps represent the zones where the behavioral syllables occur more. **(I-K)** Diagram on right: recreation of the multi-point tracking analysis sequence. Social safe, n=8; Social threat, n=9.

To determine if sub-second behavioral differences are present during social threat recognition, we used the same multi-point tracking approach as aboveto assess changes in the three-chamber social interaction assay. We first analyzed behavioral syllables during the chamber habituation period (Supplementary Figure 3), and found no differences between groups on body or tail syllables (Supplementary Figure 3D-E). When the same stimulus mouse from the learning phase was introduced into the assay, we found two different body syllables (Figure 2 I-J), and one different tail syllable, when behaviors were analyzed in this arena (Figure 2K). These behavioral syllables were exhibited in distinct chamber locations (Figure 2L-N), potentially indicating correlations with exploratory tendencies and the perceived level of threat experienced by the mice at a given location. When assessing potential distinctions between sexes across groups, we found that males in the social threat group displayed less of syllable 1 as compared to females (Supplementary Figure 5). All together, these results suggest that when mice recognize a social threat, they display unique sub-second behavioral sequences.

### Social threat recognition alters the activity of neurons in the nucleus accumbens, the paraventricular thalamus, the ventromedial hypothalamus, and the nucleus of reuniens

To investigate brain regions involved in modulating social recognition, we conducted immunohistochemistry to assess the number of cFos+ cells in mice that underwent social threat learning (or social safety learning). To accomplish this, we forced social interaction 24 hrs after social learning and extracted brains for immunohistochemistry to assess cFos+ neurons across the brain of social threat and social safe mice. In addition, we exposed a third group to the same chamber containing a non-animated object of similar size to a mouse (Figure 3A). The number of cFos+ cells in the social threat and social safe groups was normalized relative to the non-social group. Our analysis revealed significant differences in NAc, PVT, VMH, and RE (Figure 3E), whereas all other regions tested did not show statistical differences. Since previous reports have found statistical differences in mPFC using a similar task [21], we tested cFos expression across layers in an independent manner and found a significant difference in IL layers 2-3 and 5 (Figure 3E), but no difference in any of the other layer in IL, or in any of the individual layers in PL (Supplementary Figure 6). Interestingly, we identified that only layers 2-3 and 5 in IL, as well as NAc increase their activity in the social safe group (Figure 4D-E). These regions play pivotal roles in reward processing and threat extinction memory, and interact with other brain regions that are crucial for social interaction. Surprisingly, we did not find differences in PL or its layers, nor LA, BLA, CeA, MeA, or BNST (Figure 3E, Supplementary Figure 6), which have been described as regions that are pivotal for displaying defensive responses [21,51–56]. The use of a dynamic conditioned stimulus in our task likely engages multiple sensory modalities in the mouse. Unlike static stimuli, dynamic cues activate various sensory pathways simultaneously, enhancing the overall sensorial experience [57]. Studies have shown that multisensory integration occurs in the thalamus, where inputs from different senses converge. Dynamic stimuli are more effective in engaging these regions, leading to robust neural responses [58,59]. Although we did not directly observe significant differences in these structures when comparing overall cFos expression, both are intricately linked to other brain regions where we did detect significant differences (Figure 3E). It is possible that despite the absence of direct differences in these regions, they may still play a coordinated role alongside other structures that are crucial for modulating defensive responses and social interactions to facilitate social threat recognition.

**Figure 3.**
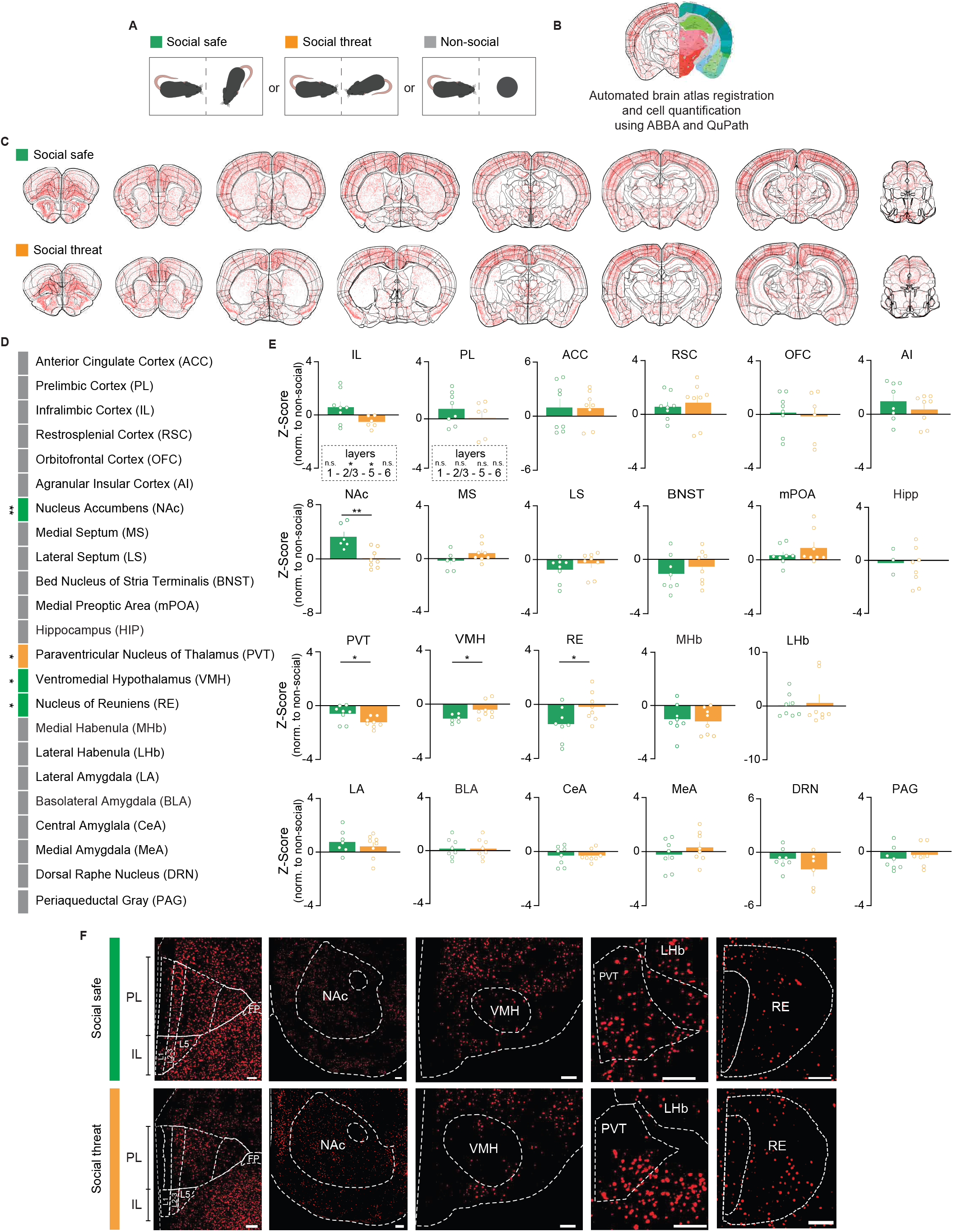
Social threat recognition induces distinct patterns of neural activity compared to social safety. **(A)** Schematic representation of the forced social interaction. **(B)** Left: Image representation of traces of cell detections for immunolabeled cFos+ cells. Right: Image taken from the interactive mouse brain atlas from the Allen Institute. (**C)** Traces of cell detections for representative cFos immunolabeled sections across conditions following atlas registration. **(D)** List of regions analyzed following cFos immunohistochemistry. The green color shows that the social safe group had the greatest change compared to the control group. The orange color shows that the threat group had the greatest change compared to the control group. The gray color shows regions that have no changes. **(E)** cFos+ cell count normalized to the non-social group shows statistical difference in IL layers 2-3 (t(12) = 2.34, p = 0.03) and 5 (t(12) = 2.92, p =0.01), NAc (t(12) = 3.80, p = 0.002), PVT (t(14) = 2.45, p = 0.02), VMH (t(12) = 2.36, p = 0.03), and RE (t(14) = 2.19, p = 0.04). All other regions did not show significant differences. **(F)** Representative micrographs showing cFos expression of social safe and social threat groups for the regions that had a significant difference. Non-social, n=8; Social safe, n=8; Social threat, n=8.

**Figure 4.**
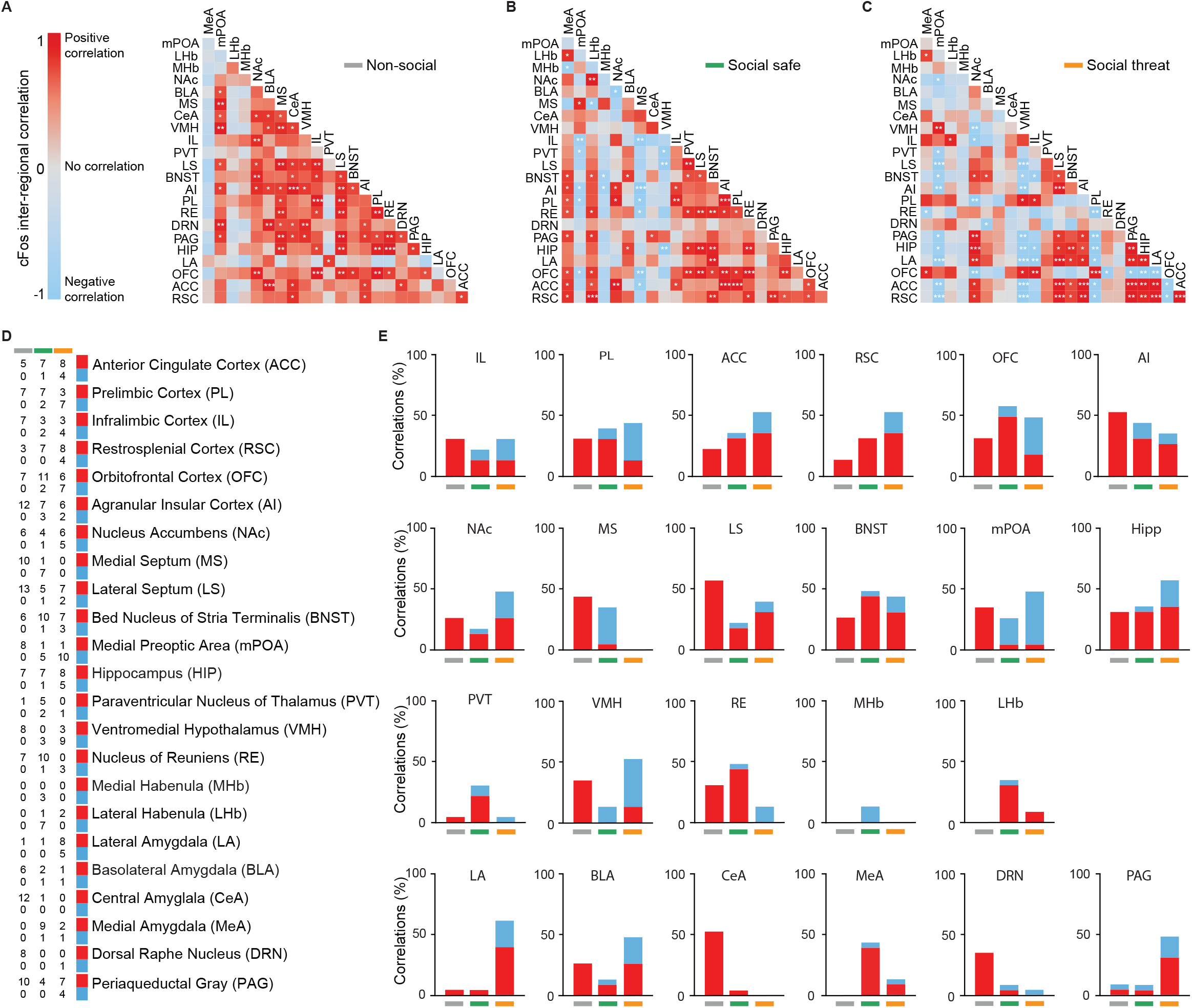
Social threat recognition reshapes functional brain connectivity. **(A-C)** Correlation of cFos cell density amongst regions calculated and plotted within a correlogram heatmap. **(D)** List of regions analyzed by cFos immunohistochemistry along with the number of significant positive (red) and negative (blue) correlations exhibited with other regions. **(E)** Percentage change in correlations across analyzed regions. Non-social, n= 8; Social safe, n=8; Social threat, n=8.

### Social threat recognition reshapes functional brain connectivity

Upon analyzing cFos expression across multiple structures, we found significant differences between social threat and social safe groups across four regions (NAc, PVT, VMH, and RE). Given the interconnected nature of these brain regions, alterations in the activity of one structure can deeply influence its communication with others [60,61]. To investigate potential shifts in inter-regional communication, we analyzed functional connectivity across these regions by computing correlations in cFos+ cell count (Figure 4). We first displayed correlation matrices of the regions for the three behavioral groups to distinguish significant positive or negative correlations that might differ between conditions (Figure 4A-C). Our analysis revealed a notable shift in correlation patterns, characterized by a drop in positive correlations and the emergence of negative correlations within social safe and social threat groups compared to the non-social group (Figure 4D-E, Supplementary Figure 7). Out of a total of 484 possible correlations, the non-social group exhibited 72 positive correlations, the social safe group displayed 55 and the social threat group showed 42. This is indicative of a drop in positive correlations when a social stimulus was present, and a further drop when it was a social threat. No negative correlations were observed in the non-social group, while the social safe and social threat groups exhibited 19 and 39 negative correlations, respectively. This shift is indicative of an emergence of negative correlations when the social stimulus was present, especially when it was a social threat. A drop in positive correlations across six structures occurred in both the social safe and social threat groups (MS, LS, mPOA, VMH, CeA and DRN), suggesting it may be related to the presence of a social stimulus. Similarly, the emergence of negative correlations was observed in mPOA and VMH across both social safe and social threat groups. Conversely, we observed positive correlations exclusively within the social safe group across six different regions (OFC, BNST, PVT, RE, LhB, MeA), possibly indicating signaling of a safe social interaction.

In contrast, we identified 11 regions where negative correlations emerged within the social threat group (ACC, PL, RSC, OFC, NAc, BNST, Hipp, VMH, LA, BLA, and PAG), and 2 regions where positive correlations emerged within the social threat group (LA and PAG). Out of the regions we previously found to be different: (1) NAc showed an emergence in negative correlations with a social threat, (2) PVT had positive correlation that emerged with the social safe and went away with the social threat, (3) VMH had a drop in positive correlations with both social safe and threat, while negative correlations were predominant with a social threat, and (4) RE showed a complete drop in positive correlations in the social threat group, as compared to the other two groups. Taken together, these 4 regions may represent important hubs for inter-regional communication across the brain to signal social safety or social threat.

## Discussion

We investigated the repertoire of behavioral responses that occur when mice learn to recognize a social safe vs a social threat stimulus, as well as how this type of recognition reshapes global brain activity. By employing social threat learning tasks with both single-point and multi-point tracking, our study revealed that an aversive social experience creates a social threat memory that evokes unique socially motivated responses the following day and reshapes functional connectivity. Specifically, we observed a drop in positive correlations and an emergence in negative correlations across brain regions in the presence of a social stimulus, that further dropped and emerged in the presence of a social threat. These findings add to a growing body of evidence showing that social recognition recruits many brain regions to facilitate appropriate behavioral responses. We believe our study provides a blueprint of behavioral sequences and brain networks that emerge to regulate socially safe and threatening encounters.

Classic tone-fear or context-fear conditioning paradigms are commonly employed to elucidate the neural processing of threats [62–67]. These models are typically associated with specific sensory modalities and related brain regions [51,56,68]. In contrast, social threat learning integrates multiple sensory inputs from tactile, olfaction, visual and auditory processing [20,69–72]. This multimodal information processing implies the engagement of many brain structures, each capable of eliciting and modulating unique defensive behaviors. In this study, utilizing single-point tracking, we observed that mice subjected to social threat learning exhibited classic defensive behaviors like freezing and avoidance. Employing unsupervised behavior clustering allowed us to extract subtle sub-second body and tail dynamics that correspond to social encounters and the experience of associating social interaction with foot-shock. It is well-documented that tail rattling is a prominent feature in rodents that is associated with aggression and territorial behavior [73–75]. Our multi-point tracking revealed six tail behaviors that are more expressed in the social safe group, with tail extension being the principal characteristic (Figure 1M-R). In the social threat group, we observed an increase in four tail behaviors exhibiting more complex movements resembling tail folding or rolling, suggesting an adaptive response to social threat presence (Figure 1S-V).

Evolutionarily, organisms often exhibit a repertoire of behaviors during initial encounters with a stimulus, but subsequently, based on previous experiences, they display a subset of these behaviors upon repeated exposure, a phenomenon termed behavioral evolution [76–78]. Upon re-exposure to the threat stimulus on the following day, we observed an increase in a syllable potentially linked to freezing behavior in the threat group, alongside a decrease in an exploratory-related syllable. Remarkably, tail movement analysis revealed that the threat group displayed a syllable increase only when approaching the social stimulus, which did not occur in the zones devoid of social stimuli (Figure 2J-K). This correlation sheds light on the contextual relevance of exhibited behaviors and their stimulus-dependent modulation.

In this study, we employed an analysis of cFos expression to identify changes in neural activity between groups across multiple brain regions (Figure 3). We noticed an increase in NAc, and layers 2/3, and 5 in the IL for the social safe group. It is well documented that the NAc plays a crucial role in social reward interactions [27,29]. Our experimental setup involved isolating mice for 24 hours before the experiment, with their only opportunity for social interaction occurring during the task. The IL is known for its involvement in both processing safety cues, and in regulating threat memories [79–85], as well as social behaviors [86,87]. Since mice prefer social interaction over isolation [10,27,88–90], the observed heightened activity in NAc and IL may be linked to the opportunity for social interaction and the presence of another mouse that was associated with safety. Conversely, we observed significant differences in activity in the VMH, PVT and RE, which are implicated in threat memory and social behaviors [20,91–95], but are not canonical threat regions [21,51,56,80,96,97] (Figure 3). On a global scale, we identified a reorganization of brain activity. We also identified that social safe and social threat experiences reshaped functional brain connectivity, by decreasing positive correlations, and increasing negative correlations (Figure 4, Supplementary Figure 6). This network analysis highlights the changes in functional connectivity across brain regions relevant for social recognition.

A notable finding was that the MeA, which exhibited no significant correlations with any structure in the group without social interaction (non-social), showed multiple correlations in the social safe group, which decreased markedly in the social threat group, as illustrated in Figure 4A-C. Recent studies have highlighted the role of MeA in modulating aggressive and mating behaviors that facilitate survival [71,72], with lesions in this region affecting such behaviors [98]. For example, MeA projections to the BNST have been found to promote aggression [99,100]. MeA also projects to the RE [91,101,102]. The RE is intricately connected to a variety of brain regions, including the PL, Hipp, PVT, and BNST, among others [91,103–106,106,107]. These connections position the RE as a pivotal area involved in the modulation of spatial context, arousal, attention, and defensive behaviors. Here, we showed that the RE exhibits a positive correlation with the MeA within the social safe group, in contrast to its negative correlation with the MeA in the social threat group (Figure 4A-B). This dichotomy underscores the potential multifunctional role the RE may play in mediating social behavioral responses. Another interesting change in functional connectivity based on the type of stimulus is within the mPOA, which, opposite to the MeA, has multiple positive correlations in the non-social group, but positive correlations dropped while negative correlations emerged strongly in the presence of a social stimulus (Figure 4 A-C). The LS is one of the most representative changes between these two groups going from a significant positive correlation with mPOA in the non-social group to significant negative correlations within both social groups (Figure 4 A-C). The LS is involved in regulating motivated behaviors [108,109], via communication with BLA [90,110,111], PL [90,108,112], NAc [90,108], VTA [108,113,114], and IL [112,113], among others. Top-down regulation throughout these brain regions show how the brain may be reshaping functional activity in order to choose the appropriate behavioral response to a salient social stimulus.

These findings collectively underscore that mice exhibit a variety of social behaviors that are dependent upon the valence of the stimulus presented. Furthermore, these experiences prompt a reorganization of functional connectivity that facilitate the most appropriate behaviors necessary for survival. Future investigations will be relevant to elucidate the neural circuits that modulate subtle behaviors and ascertain whether the manipulation of these circuits can causally regulate specific behavioral outcomes. Such studies hold promise for advancing our understanding of the neural basis of behavioral responses to social threats and may have implications for therapeutic interventions targeting maladaptive behaviors characteristic of a variety of psychiatric conditions.

## Acknowledgments

We thank Micah Baldonado and Chanjia Cai for assistance in initial key-point moseq analysis; Samir Patel and Ruben Garcia-Reyes for troubleshooting initial behavioral experiments; and Ayden Ring and Ellora McTagart for technical assistance. This work was supported by the Hellen Lyng White Fellowship (G.V.H.), Royster Fellowship (V.R.C.), the National Institute of Neurological Disorders and Stroke (UNC Neuroscience Curriculum: T32 NS007431, N.W.M.), the National Institute of General Medical Sciences (UNC PREP: R25 GM089569, C.M.R.-P.), the UNC Department of Applied Physical Sciences (N.C.P.), the Burroughs Wellcome Fund (N.C.P.), a Beckman Young Investigator Award (N.C.P.), the UT Program on Science and Technology Acquisition and Retention (STARs, A.B.R.), the UTSA College of Sciences (A.B.R.), the National Institute of Child Health and Human Dvelopment (IDDRC; P50 HD103573; PI Gabriel Dichter), the UNC Department of Psychiatry (J.R.-R.), the Foundation of Hope (A.S.Z. & J.R.-R.), the NC TraCS Institute (J.R.-R. & N.C.P.), the Brain and Behavior Research Foundation (J.R.-R), the Kavli Foundation (J.R.-R.), The Whitehall Foundation (J.R.-R.), and the National Institute of Mental Health (R01 MH132073, J.R.-R).

## Author Contributions

Conceptualization: J.R.-R., A.B.R., A.S.Z. Research Designs: J.R.-R., A.B.R., N.C.P., S.S.M., G.V.H., N.W.M. Methodology: J.R.-R, G.V.H., N.W.M., V.R.C. Behavioral Experiments: G.V.H., N.W.M. Data Analysis: G.V.H., N.W.M., V.R.C. Histology: N.W.M., C.M.R.-P., S.M.L. Writing Article: G.V.H., N.W.M., V.R.C., J.R.-R., A.B.R. Supervision of all Aspects of the Study: J.R.-R.

## Data Availability

All data analysis scripts and raw data supporting this research will be shared for appropriate scientific use upon request.

## Competing Financial Interests

The authors declare no competing financial interests.

## Notes

### Competing Interest Statement

The authors have declared no competing interest.

