## Supplemental Material for "Social threat alters the behavioral structure of social motivation and reshapes functional brain connectivity"

**Supplementary Methods for Velazquez-Hernandez et al., Social Threat**

Subjects. Prior to the experiment, the experimental mice were group-housed until 24 hours before the initiation of social threat learning, at which point they were transitioned to individual housing. Stimulus mice with lower body weight, same-sex, and comparable age to the experimental animal were selected. The stimulus mice were group-housed during the entire experiment. All animals were maintained on a reverse 12-hour light-dark cycle (lights off at 08:00 AM) with *ad libitum* access to food and water. Behavior was tested during the dark cycle.

Social threat learning. Our test chambers were enclosed within a custom sound-attenuating box measuring 81x86 cm, to reduce any external noise and visual stimulation that might affect the behavior of the mice. The floor of the modular test chambers consisted of stainless-steel bars that deliver electric foot-shocks through a computer interface (ANY-maze; Stoelting Co., IL, USA). We replaced one of the metal panels of the modular test chambers with a transparent plexiglass panel containing four windows (0.8 cm. width, 2.2 cm tall) to allow both, experimental and stimulus mice, to interact through sniffing, nose pokes, and sight.

Stimulus mice were habituated to the social chambers for 3 days, 10 minutes each day. Experiments began by placing the experimental mouse in the modular test chamber for a 10-minute adaptation period. To avoid the stress that may be associated with the initial introduction of the stimulus mouse, a barrier was placed in front of the social interaction window. Following the placement of the stimulus mouse, the wall was removed and the mice were allowed to interact for 10 min. Mice that never approached the social stimulus during the session or that exhibited freezing above 40% during the initial two minutes (prior to any social contact) were excluded from the experimental cohort (n=2). All mice were returned to their home-cage after 10 minutes period of social interaction.

The arena was divided into three sections (20.5 cm for the ones on the side and 16 cm for the neutral zone) by opaque walls with rectangular openings (9.5 cm wide), all uniformly illuminated with white light (54 lux). To house the stimulus mouse, we used triangular social boxes meant to fit into the corners of the three-chamber assay (10.5x10.5x15.3 cm, 25.5 cm high). These social boxes featured holes (1 cm) instead of the rectangular windows employed during social threat learning to eliminate potential contextual cues.

At the beginning of the trial, experimental mice were initially allowed to explore the three-chamber assay for 10 minutes to habituate them to the context. Following this adaptation period, the mouse was gently relocated to the central chamber, and two opaque walls were added to occlude the rectangular openings. We then placed the social boxes on either side of the arena and introduced the stimulus mouse on the side that the experimental mouse explored more during the initial 10-minute adaptation period. Subsequently, the opaque walls were removed, enabling the experimental mouse to explore the entire arena for an additional 10 minutes.

Three measures of defensive responses were assessed throughout behavioral experiments using one-point tracking data: 1) percent of time spent freezing, 2) time in the social zone, and 3) social preference. Freezing was defined as the absence of all movement except respiration. The amount of time freezing during the test was expressed as the percentage of the test duration. Time in the social zone was defined as the amount of time that the experimental mice spent in the social area. The amount of time in the social zone during the test was expressed as the percentage of the test duration. Social preference was defined by comparing time in the social zone ($t_{social}$) and the time in the nonsocial zone ($t_{non\text{-}social}$) with the following formula:

$$\frac{t_{social} - t_{nonsocial}}{t_{social} + t_{nonsocial}}$$

All behavioral responses were recorded with digital video cameras (Imaging SourceIMAGING SOURCE model DMK22BUC03, Charlotte, USA) and automatically analyzed with commercial software using the centroid of the mouse to define its location (ANY-maze; Stoelting Co., IL, USA).

<u>Multi-point tracking and motion sequencing.</u> The model was allowed to train for 100,000 iterations with the loss reduction plateauing at 0.002. To account for the temporal distribution of differences

between sub-millisecond body and tail-based behaviors, we trained a separate model for body and tail keypoints. The MoSeq model trained with tail-based key points converged to 4 frames, while the body-based model converged to 6 frames, indicating the median frame duration across behaviors (for videos captured at 15 frames per second, this indicates a median frame duration of ~267 and 400 ms respectively).

Immunohistochemistry. Mice were perfused with phosphate buffered saline (PBS) (*Sigma-Aldrich* cat #P4417), and paraformaldehyde (PFA, 4%) (*Sigma-Aldrich* cat #158127). Brains were collected and post-fixed in 4% PFA. 24 hours later, the brains were transferred into a 30% sucrose (Fisher cat #S5-3) in PBS solution. Brains were sectioned at 40 μm using a cryostat (Leica, CM 3050 S) at −20 °C. Sections were collected into tissue culture plates with PBS. On the first day of immunohistochemistry, the sections were washed 3x in PBS for five minutes on a shaker at room temperature. Then, the sections were transferred to a blocking solution (Normal Donkey Serum 5% + 0.2% Triton X-100 in PBS) for 1 hr. Tissue was then transferred to the primary antibody solution (Rabbit anti-cFos (Synaptic Systems 226-008) 1:5000, 5% NDS + 0.2% Triton X-100 in PBS) and were incubated overnight on a shaker at 4 °C. On the second immunohistochemistry day, sections were washed three times in PBST (0.2% Triton X-100 in PBS) for five minutes on a shaker at room temperature. Then, the sections were transferred to the secondary antibody solution (Donkey anti-Rabbit Alexa Fluor 647 (Invitrogen A311573) 1:2000) to be incubated for 1 hr. on a shaker three times for five minutes in PBST at room temperature. After secondary incubation, the sections were washed three times with PBS. Sections were mounted onto charged slides with Fluromount-G with DAPI (*Southern Biotech* cat # 0100-20) mounting medium and coverslipped. Damaged tissue sections were excluded from further analysis.

Cell counting and analysis. cFos+ nuclei were quantified with special attention in the anterior cingulate cortex (ACC), prelimbic (PL) and infralimbic cortex (IL), retrosplenial cortex (RSC), orbitofrontal cortex (OFC), anterior insular cortex (AI), nucleus accumbens (NAC), medial (MS) and lateral septum (LS), bed nucleus of stria terminalis (BNST), medial preoptic area (mPOA), hippocampus (Hip), paraventricular nucleus of thalamus (PVT), ventromedial hypothalamus

(VMH), nucleus of reuniens (RE), medial (MHb) and lateral habenula (LHb), lateral (LA), basolateral (BLA), central (CeA), and medial amygdala (MeA), dorsal raphe nucleus (DRN), and periaqueductal gray (PAG). All images for cFos analysis were taken with a fluorescence microscope (Keyence BZ-X810, IL, USA) with a 20x objective (0.75NA, 0.9WD).

### Supplementary Figures

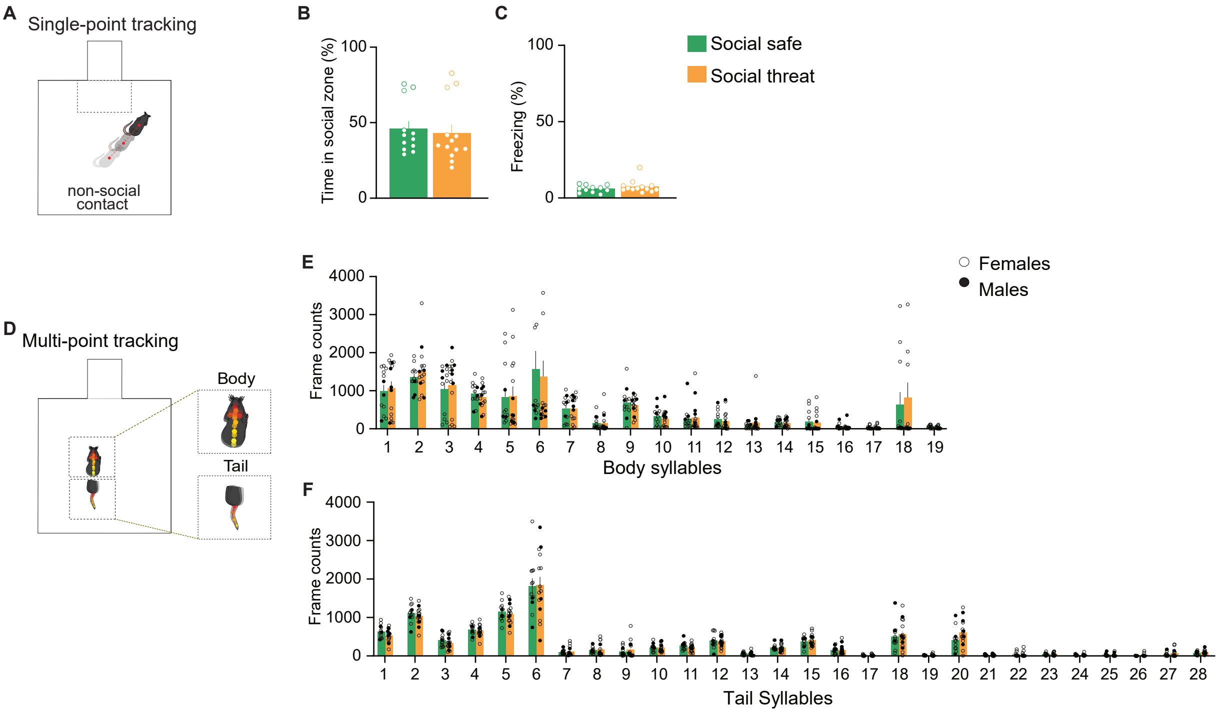


**Supplemental Figure 1. Baseline Assessment on day 1 Using Single-point and Multipoint Tracking.** We examined mouse behavior during a 10-minute adaptation period on day 1 using single-point tracking **(A)** to measure time spent in the social zone **(B)** and freezing behavior **(C)**. Additionally, we utilized multipoint tracking **(D)** to analyze behavioral syllables related to body **(E)** and tail **(F)**. Our analysis revealed no differences between groups in either single-point or multipoint tracking analyses.

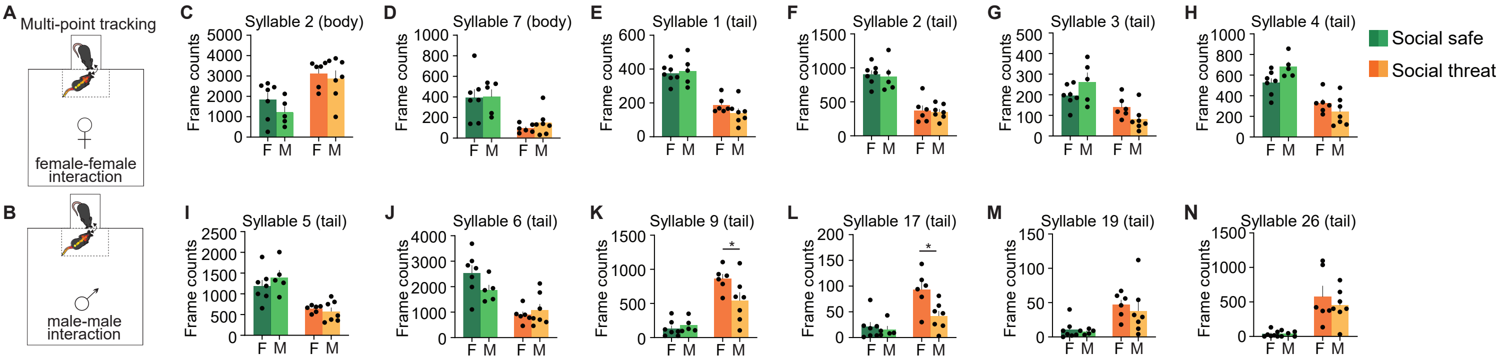

**Supplementary Figure 2. Sex difference in syllables from body and tail across groups on day 1. (A-B)** Comparison between female and male mice in both experimental conditions to test any differences in the syllables on body and tail, exhibiting significant differences. **(C-D)** No differences between sexes across syllables in body. **(E-N)** No differences between sexes across syllables in tail in the social safe group. **(K)** Females in the social threat group exhibited an increase in activity in syllables 9 ($F(3, 15) = 19.84$, $p = 0.0001$, posthoc comparison $p = 0.01$) and 17 from the tail ($F(3, 15) = 9.03$, $p = 0.001$, posthoc comparison $p = 0.03$. (social safe, $n = 12$; social threat, $n = 13$).

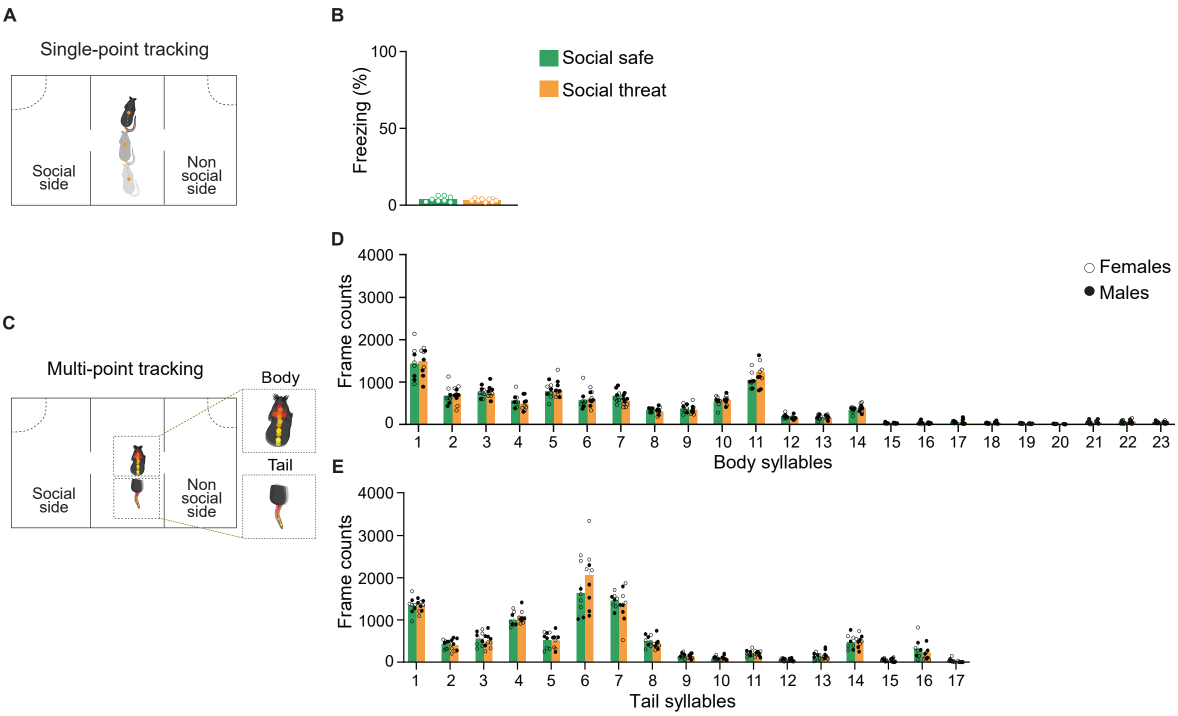


**Supplementary Figure 3. Baseline assessment on day 2 using single-point and multi-point tracking. (A) Representation of single-point tracking to analyze freezing behavior. (B)** Single-point analysis shows no difference between groups on freezing analysis. **(C)** Representation of multi-point tracking to analyze behavior syllables related to body and tail. **(D)** Multi-point tracking analysis revealed no differences between groups for the body syllables. **(E)** Multi-point tracking analysis revealed no differences for tail syllables. (social safe, n= 8; social threat, n= 9)

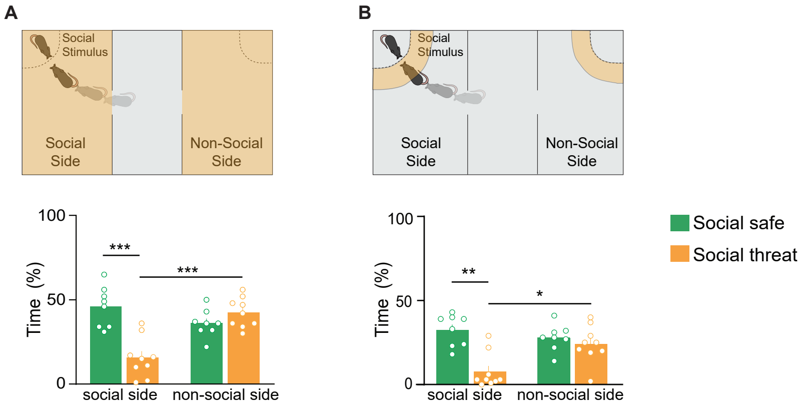


**Supplementary Figure 4. Social threat conditioning induces social avoidance. (A)** Top: Schematic diagram illustration of the three-chamber social interaction assay. Bottom:The social threat group exhibited reduced time spent on the social side and increased time spent on the non-social side ($F(3, 19) = 11.41$, $p = 0.0002$, posthoc comparison time in social side: social safe vs social threat $p = 0.0005$. social threat group: time in social side vs time in non-social side $p = 0.0005$). **(B)** Top: Schematic diagram illustration of the three-chamber social interaction assay. Bottom: The social threat group showed decreased time spent in the social interaction zone and increased time spent in the non-social side ($F(3, 19) = 8.12$, $p = 0.001$, posthoc comparison time in the social interaction area: social safe vs social threat $p = 0.001$, social threat group: time social interaction area vs time in non-social interaction area $p = 0.01$). Transparent orange highlights the areas analyzed using single-point tracking. Social safe, n=8; Social threat, n=9.

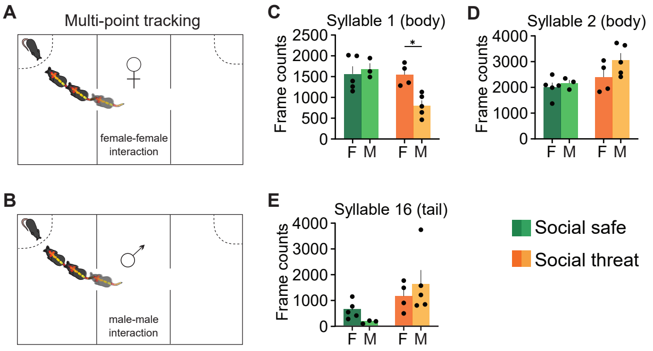


**Supplementary Figure 5. Sex difference in syllables from body and tail across groups on day 2. (A-B)** Comparison between female and male mice in both experimental conditions to test any differences in the syllables exhibiting significant differences. **(C)** Males in the social threat group exhibited a decrease in the activity in body syllable 1 ($F(3,9) = 7.94$, $p = 0.006$, posthoc comparison social threat group: females vs males $p = 0.01$). **(D)** No differences between sexes in the activity in body syllable 2 ($F(3,9) = 3.31$, $p = 0.07$), (E) or tail syllable 16 ($F(3,9) = 3.42$, $p = 0.06$). Social safe, n=8; Social threat, n=9.

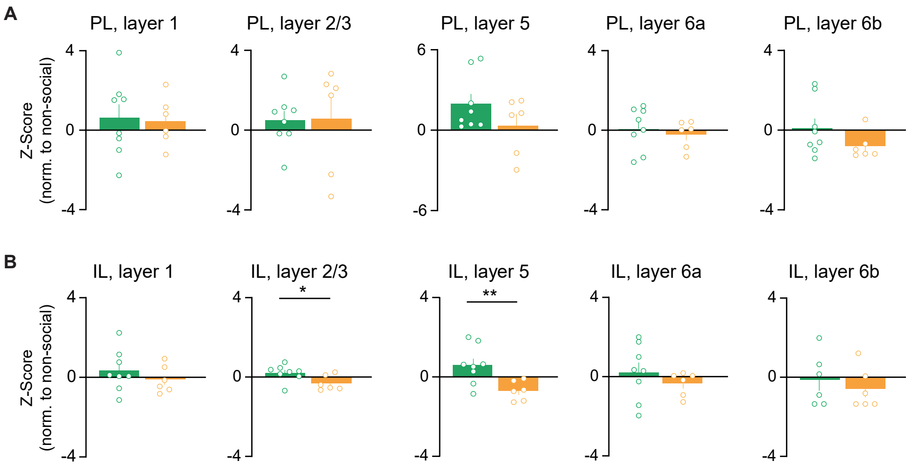


**Supplementary Figure 6. Social threat recognition induces changes in cFos expression within specific layers of the medial prefrontal cortex. (A)** cFos+ cell count in different layers of PL, normalized to the non-social group. **(B)** cFos+ cell count in different layers of IL shows significant differences in layers 2/3 ($t_{(12)} = 2.34$, p = 0.03) and 5 ($t_{(12)} = 2.92$, p =0.01), normalized to the non-social group. Non-social, n=8; Social safe, n=8; Social threat, n=6.

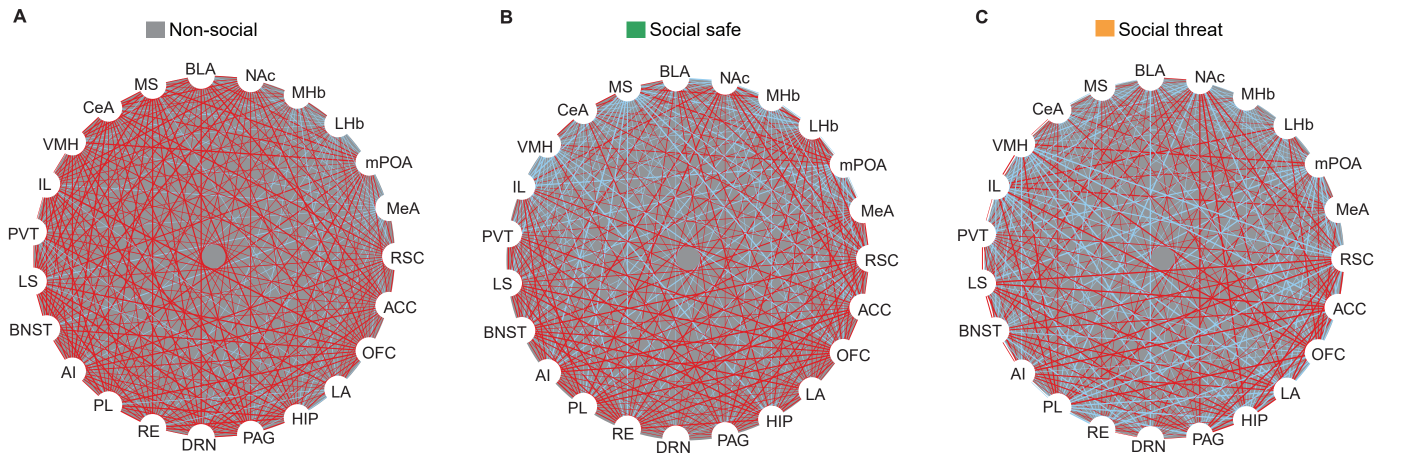


**Supplementary Figure 7. Network plots of functional connectivity using cFos analysis. (A)** Analysis of the non-social group shows strong positive correlations in cFos cell count across regions. **(B)** Analysis of the social safe group shows a decrease in positive correlations with some increase in negative correlations. **(C)** Analysis of the social threat demonstrates a notable reversal between positive and negative correlations. Red lines show positive correlations between regions. Blue lines show negative correlations between regions. Non-social, n=8; Social safe, n=8; Social threat, n=8.
